# Binding site plasticity regulation of the FimH catch-bond mechanism

**DOI:** 10.1101/2022.11.15.516604

**Authors:** Olivier Languin–Cattoën, Fabio Sterpone, Guillaume Stirnemann

**Affiliations:** CNRS Laboratoire de Biochimie Théorique, Institut de Biologie Physico-Chimique, PSL University, Université Paris Cité, 13 rue Pierre et Marie Curie, 75005, Paris, France

## Abstract

The bacterial fimbrial adhesin FimH is a remarkable and well-studied catch-bond protein found at the tip of *E. coli* type 1 pili, which allows pathogenic strains involved in urinary tract infections to bind high-mannose glycans exposed on human epithelia. The catch-bond behavior of FimH, where the strength of the interaction increases when a force is applied to separate the two partners, enables the bacteria to resist clearance when they are subjected to shear forces induced by urine flow. Two decades of experimental studies performed at the single-molecule level, as well as X-ray crystallography and modeling studies, have led to a consensus picture whereby force separates the binding domain from an inhibitor domain, effectively triggering an allosteric conformational change in the former. This force-induced allostery is thought to be responsible for an increased binding affinity at the core of the catch-bond mechanism. However, some important questions remain, the most challenging one being that the crystal structures corresponding to these two allosteric states show almost superimposable binding-site geometries, which questions the molecular origin for the large difference in affinity. Using molecular dynamics with a combination of enhanced-sampling techniques, we demonstrate that the static picture provided by the crystal structures conceals a variety of binding-site conformations that have a key impact on the apparent affinity. Crucially, the respective populations in each of these conformations are very different between the two allosteric states of the binding domain, which can then be related to experimental affinity measurements. We also evidence a previously unappreciated but important effect: in addition to the well-established role of the force as an allosteric regulator via domain separation, application of force tends to directly favor the high-affinity binding-site conformations. We hypothesize that this additional *local* catch-bond effect could delay unbinding between the bacteria and the host cell before the *global* allosteric transition occurs, as well as stabilizing the complex even more once in the high-affinity allosteric state.

## Introduction

Catch bonds form a peculiar family of non-covalent molecular bonds that display a counter-intuitive behavior: their lifetime is increased under mechanical tension – much in contrast to traditional bonds (or slip bonds), whose lifetime often decreases exponentially with tensile force.^1–4^ Catch-bond properties have attracted growing interest in recent years due to their pervasive involvement in mechanobiological phenomena, including cell-cell and cell-matrix interactions, blood coagulation, cellular motility, and pathogen-host adhesion.^4–11^

While catch bonds have been conceptually proposed a few decades ago,^1^ characterization and rationalization of their properties was only achieved in recent years, in particular thanks to the development of experimental techniques that enabled to replicate in a laboratory set-up the microscopic forces that operate in vivo, either at a single-cell or at a single-molecule level.^12–14^

The bacterial fimbrial adhesin FimH is a remarkable and well-studied catch-bond system found at the tip of *E. coli* type I pili.^15^ It consists of two domains: a pilin domain connected to the rest of the pilus, and an apical lectin domain carrying a binding site specific to mannose derivatives and mannose-terminated glycans expressed at the surface of urothelial cells.^16^ This allows the bacteria to anchor human epithelia in the lower urinary tract, possibly leading to urinary tract infections (UTIs).^17,18^ UTIs are one of the most common bacterial infections – impacting almost one in two women in their lifetimes –, and one of the major causes of antibiotic prescription. As such, rising concerns regarding multidrug resistance have prompted researchers to investigate other alternatives, such as anti-adhesion therapies that target the binding site of FimH or the pilus assembly pathway. ^18–21^ However, it has been widely acknowledged that the catch-bond behavior of FimH enables the bacteria to resist clearance when they are subjected to shear forces induced by urine flow, while retaining their mobility in static conditions and being less sensitive to soluble decoys. ^20^ Hence, although anti-adhesion therapies based on competitive inhibitors targeting FimH may be good candidates as a complement or replacement to antibiotics, their efficiency is likely to be impaired by the target’s catch-bond properties. A thorough understanding of the detailed mechanism of the catch bond behavior of FimH is thus necessary for devising appropriate anti-adhesive strategies.

The state-of-the-art picture of FimH’s catch-bond machinery can be understood as force-induced, negative intramolecular allostery, whereby mechanical strain on the protein-ligand complex induces coaxial separation of the two protein domains and extension of the inter-domain linker, followed by an allosteric transition of the binding domain to a higher affinity state (Fig. 1). This global framework has emerged from kinetic and thermodynamic experiments, directed mutagenesis experiments, and a vast body of crystallographic (and a few solution) structures obtained in various conditions.^22–29^ Pioneering work from Thomas et al. provided results that fitted a “two-state” (that is, two *bound* states and one *unbound* state) kinetic model based on parallel-plate flow chamber experiments.^24^ However, a proper structural evidence of the protein allosteric mechanism has long remained difficult to obtain because purified FimH is not stable and prone to aggregation. For that reason, early crystal structures focused on the isolated lectin domain and did not portray the full protein in physiological-like conditions.

**Figure 1:**
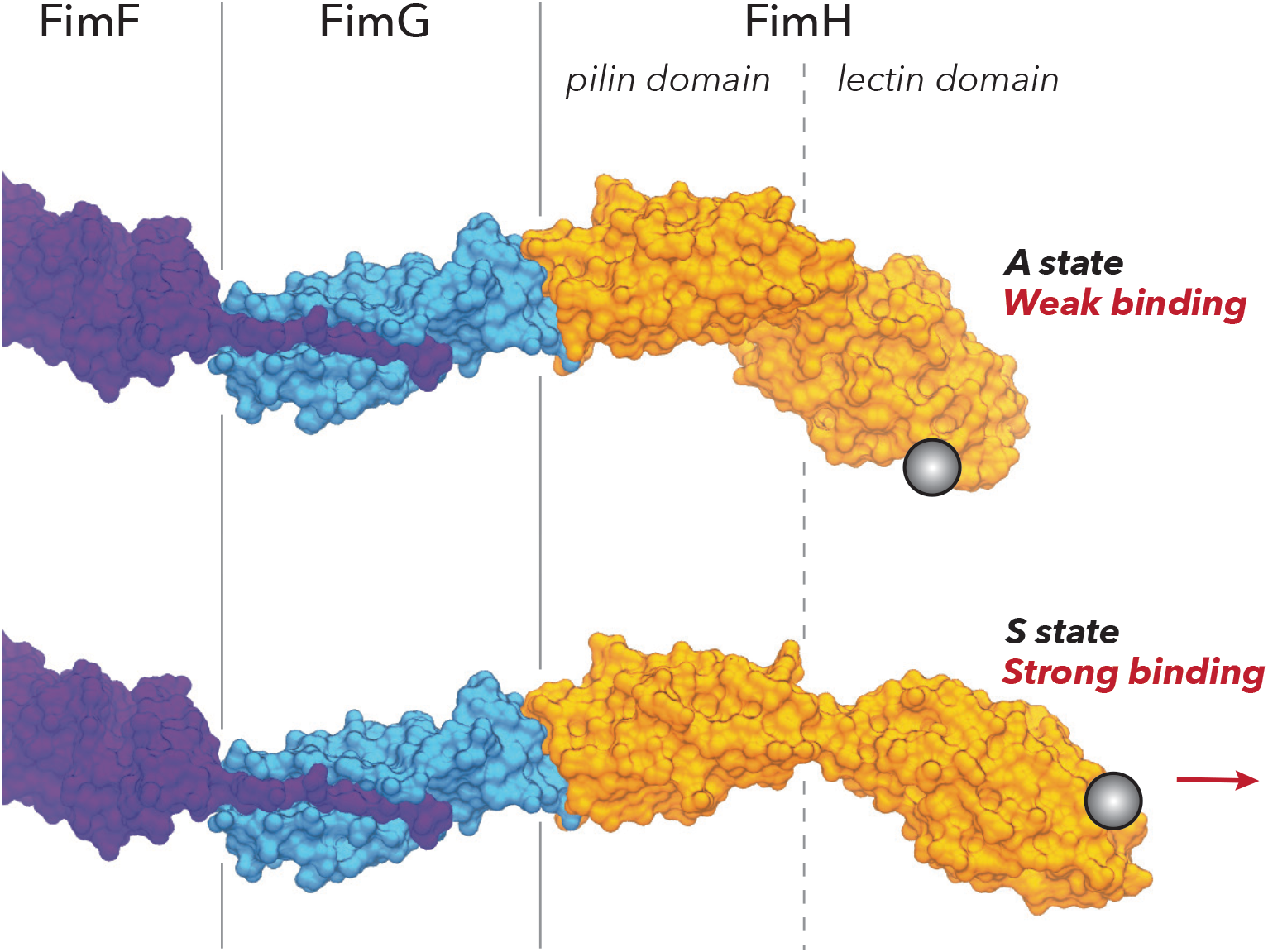
General mechanism of the FimH catch bond. FimH *(orange)* is located at the pilial extremity and connected to the rest of the pilus through stable *donnor-strand complementation* to the FimG *(blue)* and FimF *(violet)* pilin units. FimH’s lectin domain comprises a mannose-specific binding site able to anchor the bacteria to epithelial proteoglycans *(gray circle)*. When the bond is subjected to mechanical tension, the bound lectin domain separates from the pilin regulatory domain and undergoes allosteric transition from a “weak” to a “strong” binding state. *Composite models based on PDB* 3jwn^*22*^ *and* 4xoe.^*23*^

In vivo, the pilin domain is stabilized by the non-covalent binding of a beta strand from the following unit of the pilus apex, FimG. This process, coined “donor-strand complementation”, is also responsible for the assembly of the remaining pilial units (FimF and poly-FimA) and ensures the exceptional solidity of the fimbria^30,31^. Eventually, Sauer et al. could stabilize FimH with the appropriate 14-aminoacid “donor strand” peptide (DsG) and they carried out kinetic and equilibrium measurements as well as X-ray crystallography. ^23^ They found that the isolated lectin domain (that we will hereafter denote by *L* for simplicity) shows a binding affinity several thousand times higher than the full protein construct (FimH·DsG, that we will write *LP*), thus demonstrating the inhibitory action of the pilin domain. They proposed a thermodynamic cycle made of four states (two *bound* states and two *unbound* states) and, for the first time, tried to associate the thermodynamic states of the model to structural features of the crystal structures.^23,32^

However, there is no straightforward correspondence between the states of a given kinetic or thermodynamic model (derived to explain experimental measurements) and the structures obtained through crystallography or NMR. The native crystal structure may not be fully representative of the biologically-active state, ^33,34^ and often hides important dynamical fluctuations and conformational changes that are very relevant for the protein function (While B-factor analysis provides valuable insight on plasticity, there is some evidence that traditional refinement processes tend to underestimate protein conformational heterogeneity^35^). In the case of FimH, crystal structures of the ligand-bound lectin domain in presence or absence of the pilin domain show remarkably similar binding site geometries. ^23^ This contrasts with the very large difference in unbinding rates measured for the corresponding systems in solution. Indeed, the sheer difference in *K*_*D*_, going from 3.0 × 10^−9^ m for *L* to 9.9 × 10^−6^ m for *LP*, is mainly due to strikingly divergent unbinding rates (*k*_off_), respectively 3.5 × 10^−4^ s^*-*1^ and 5.8 ×10^1^ s^*-*1^.^23,32^ Meanwhile, the apo crystal shows a notably different binding site structure that could be a good candidate for such a low-affinity state. This calls into question the induced-fit picture that directly emerges from the apo/holo comparison.

In biophysical studies, molecular dynamics (MD) simulations appear as an ideal tool to connect the structural and kinetic/thermodynamic pictures, in the sense that they enable to probe the plasticity of relevant protein states, to investigate the collective variables involved in transitions between them, and to provide a molecular view of how a force acts on the protein conformational landscape. In the specific case of catch bonds, although various theoretical models are consistent with experimental measurements, they are often hard to discriminate using only macroscopic observables, such as kinetic and thermodynamic data, and a full understanding often benefits from molecular modeling and simulations.^2^ In particular, MD has proven useful in a number of contexts for studying force-induced changes in biomolecules and their assemblies, with implications for both the interpretation of single-molecule force-spectroscopy results^36–38^ and the understanding of mechanosensing and mechanotransduction.^39^ A number of previous studies employed such approaches for the FimH system,^23,27,30,40–44^ often in combination with experiments, but they were mostly limited to traditional, nano-to microsecond-timescale MD simulations that can hardly provide an extensive assessment of the protein conformational landscape.

In this paper, we aim to investigate how the structural plasticity of FimH dictates its catch-bond adhesive properties beyond the well-established, but limited, crystallographic evidence. To this end, we employ a combination of enhanced-sampling MD approaches to explore and to quantify the protein’s conformational diversity. We provide unprecedented assessment of the binding domain’s free-energy landscape along relevant collective variables, and quantify how it is influenced by ligand binding and mechanical strain. Together, our results suggest that the FimH catch-bond mechanism is governed by a significant population shift in the binding-site opening conformations, caused by both the allosteric transition between low and high affinity states, and also by the direct application of a tensile force on a given allosteric state. The picture that emerges from our study also explains the vastly different affinity of the two allosteric states for mannosides, despite very similar active site geometries in the crystal structures.

## Methods

### Structures, nomenclature

FimH displays notable conformational diversity, found among more than 60 structures published in the Protein Data Bank (PDB), and ranging from the lectin domain alone to the full fimbrial tip assembly, either in presence or absence of mannose-derivated ligands. As shown in a recent work based on a pairwise RMSD analysis from Magala et al.,^45^ most of the intra-domain plasticity is found in the binding (lectin) domain and can be clustered in a set of five conformers. These conformers have been inconsistently labeled in the literature, using adjectives alternatively referring to the binding-site conformation (“open” or “closed”, “loose” or “tight”, “wide” or “narrow”), the lectin-domain overall shape (“elongated” or “compressed”), the interaction with the pilin domain (“Associated (A)” or “Separated (S)”), the presence of a ligand (“free” or “bound”) or the inferred binding affinity corresponding to that state (“low affinity” or “high affinity”, “relaxed (R)” or “tense (T)” – by analogy with traditional allosteric nomenclature). The pilin domain, on the other hand, exhibits very little structural diversity. Interdomain plasticity consists of one well-defined *Associated* conformation, where the two domains are in close interaction, and an ensemble of nonspecific poses where the domains have little to no interaction and orient themselves freely thanks to the flexibility of the linker. ^28,45^

Fig. 2 presents the three main states of the lectin domain that are relevant to fimbrial adhesion and will serve as the basis to our study. They correspond respectively to groups 1, 4 and 3 in Magala et al. ^45^ (group 2 comes from a single NMR study and is nearly identical to group 1, and group 5 corresponds to a structure of the pilial tip in complex with the usher FimD,^46^ thus mostly relevant to pilial assembly pathway in the outer membrane):

**Figure 2:**
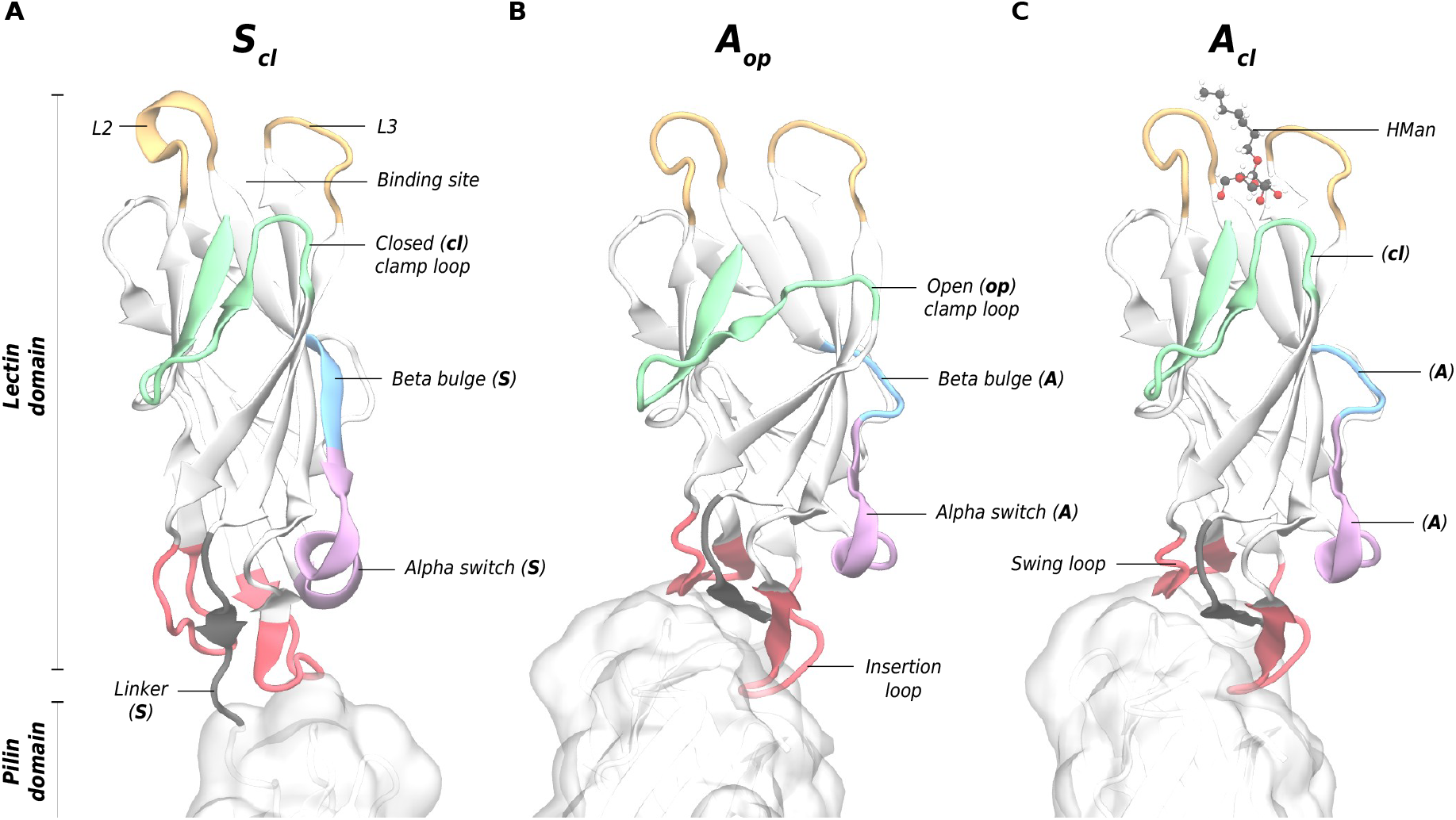
Conformational repertoire of the lectin domain. (A) The *Separated* (*S*) allosteric state of the lectin domain is characterized by a succession of a beta strand (blue) and an alpha helix (mauve) in the 59–71 region. Meanwhile, (B) and (C) the *Associated* (*A*) state presents in the same segment a bulge followed by a 3_10_ helix. The clamp loop (green) bordering the binding pocket can either be in a *closed* (*cl*) or *open* (*op*) configuration. In the PDB, *S* state always goes with a *closed* binding pocket while *A* can either be *open* (B) or *closed* (C) depending on the presence or absence of a ligand (here *α*-D-heptylmannose, HMan).

1. The first state corresponds to structures where the lectin domain – either liganted or not – is *Separated* (*S*) from the regulatory pilin domain, either because the former was crystalized in isolation, or because tight association was prevented by a third interacting partner (for example the chaperone FimC^47^) or specific mutations. Its binding site is in a *closed* (*cl*) configuration where a so-called clamp loop completes a well-defined hydrophobic ridge around the mannoside pocket. Given the high affinity of the isolated lectin domain in solution, this conformation is believed to correspond to a high-affinity state. We note this conformation *S*_*cl*_.
2. The second one corresponds to the unliganted protein where the domains are closely interacting in an *Associated* (*A*) fashion. Key secondary structure differences are observed in the lectin fold, especially in the beta-bulge/alpha-switch (residues 59-71) segment. Additionally, the binding site is found in an *open* conformation where polar, mannose-specific sidechains within the pocket are more exposed to the solvent. We label this state *A*_*op*_.
3. The third one is very similar to the second one but corresponds to cocrystals with a mannose derivative and differs in its *closed* binding site – we hence refer to this conformation as *A*_*cl*_. This has led to hypothesize an induced-fit mechanism upon ligand fixation.^23^

We emphasize here the importance of distinguishing the lectin domain’s conformational state and the composition (ligand-wise and pilin-domain-wise) of a given system, since they are in principle orthogonal. In the following we will always denote a given state using a plain letter for the allosteric form of the lectin domain (*A* or *S*) and a subscript for the opening degree of the binding site (*op* or *cl*). When needed, we will additionally specify in superscript the presence or absence of the ligand using a plain or hollow circle (• or ○), and the presence or absence of the pilin domain (*L* for lectin only, *LP* for the full FimH·DsG construct). As an example, 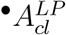 stands for the full, domain-associated protein, in presence of the HMan ligand and with a *closed* binding site.

### System preparation

Simulations were prepared from PDB structures using the GROMACS 2018.4^48^ `pdb2gmx` tool and the included AMBER ff99SB-ILDN force field.^49^ Protonation state was predicted at neutral pH using the H++ server^50^ and differed from the default `pdb2gmx` parameters only for histidine 45 (estimated p*K*_*a*_ ≈ 8–9), which we manually set to its diprotonated form. The topology for the ligand was prepared using the Generalized Amber Force Field (GAFF)^51^ using the ACPYPE^52^ tool. Systems were solvated with TIP3P water in either cubic or orthorhombic boxes ensuring at least 1.2 nm margins around the solute. Sodium and chloride ions were added at about 50 mM so as to neutralize the systems. Energy minimization and a short equilibration (200 ps) with restrained heavy-atoms positions were carried out before production runs.

### DBC Restraint

In order to prevent the ligand from escaping the binding site during the exploration of the conformational space of FimH, a Distance-from-Bound-Configuration (DBC) collective variable^53^ is defined (and later restrained) as the RMSD of (a set of) the ligand’s atoms to a reference configuration, in the binding site’s frame of reference. Let **x**_*R*_ and **x**ℓ be the coordinates associated to representative atoms of the receptor and the ligand, 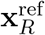 and 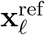 those coordinates in a reference structure (here PDB 4xoe, 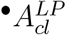 state). We note 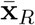 the center of mass of **x**_*R*_. After some time, the receptor will have undergone *(i)* a global translation 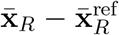 of its center of mass combined to *(ii)* a global rotation **R** around its center of mass and *(iii)* internal conformational changes. The coordinates 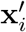 of some ligand atom *i* after correcting for the binding site roto-translational diffusion are

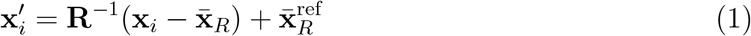

and the DBC is expressed as

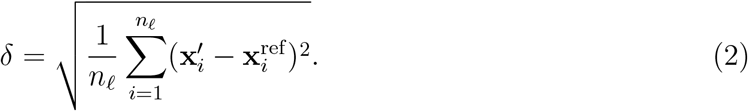

where the sum runs over the *n*ℓ representative ligand atoms. We used the mannose cycle’s heavy atoms (C1…C6, O1…O6) for the ligand set and the alpha carbon atoms of residues Phe1, Asn46, Asp54, Tyr95, Asn135 for the binding site set. We define a restraining potential to keep the ligand in a pose close to a *bound* reference, using a semi-harmonic flat-well potential:

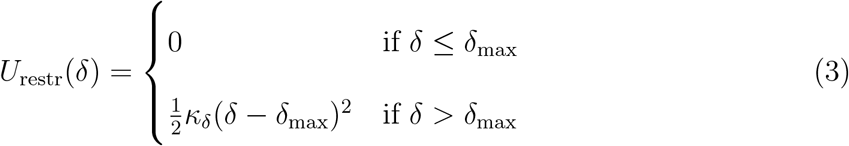

where *δ*_max_ =2.5 Å is chosen to allow comfortable fluctuations of the DBC near the bound state during REST2 sampling, as estimated using short unbiased MD trajectories (see SI). A tighter value of 1.8 Å was used during the REUS free-energy profile calculations. The harmonic constant *κ* _*δ*_ is set to 500 kJ mol^*-*1^ Å^*-*1^. The DBC and flat-well restraint potential were implemented using the Colvars^54^ module for GROMACS.

### Rotational restraints

To allow the use of a rectangular simulation box for the rather elongated two-domain systems (*LP*), and reduce the simulated solvent volume, we prevent rotational diffusion with another restraining bias *U*_rot_. The rotational component of the best RMSD alignment to a reference structure (using *α*-carbons only) is expressed as a quaternion **q** = (*q*_1_, *q*_2_, *q*_3_, *q*_4_) and restrained with a harmonic bias to the identity **q**^*I*^ = (1,0,0,0):

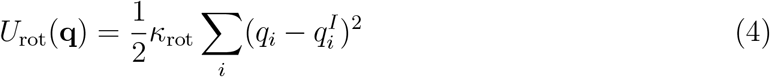

where *κ*_rot_ is set to 500 kJ mol^*-*1^. Importantly, such bias does not influence the thermodynamics of the protein internal degrees of freedom. It was implemented using the `orientation` component of the Colvars module.

### REST2

Replica Exchange with Solute Tempering (REST2)^55^ simulations were carried out using GROMACS 2018.4 patched with the PLUMED 2.5^56,57^ and Colvars modules for restraints (DBC and orientation). REST2 is a type of Hamiltonian replica exchange where the degrees of freedom of the region of interest (in our case, the lectin domain) are accelerated by downscaling torsional and electrostatic potential energy terms. REST2 improves sampling by using parallel simulations of the same system on a ladder of rescaled potentials that allow easier crossing of kinetic barriers, akin to traditional temperature replica exchange. Each replica samples its own isothermal-isobaric ensemble but may exchange configurations with adjacent replicas using a Metropolis-Hastings criterion.

For each system, a set of *N* = 20 replicas was constructed. Topologies were produced using the `partial_tempering` program of the PLUMED package. The scaling factors λ_*k*_ were chosen to obtain a range of effective temperatures between *T*_0_ = 300 K and *T*_*max*_ = 600 K following a geometric sequence

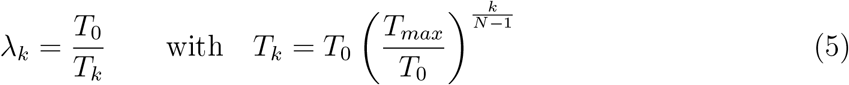

We ensured that this choice of *N* and *T*_*max*_ resulted in satisfactory exchange probabilities (0.1–0.2) between adjacent replicas.

Each system was simulated for 500 ns with exchange between replicas attempted every 2 ps. The solute configuration was recorded every 20 ps for analysis. As our first approach is only exploratory and do not aim at a quantitative assessment, we keep the full trajectory for the analysis despite the initial out-of-equilibrium “heating” phase of replica exchange methods.

### Definition of the bound state

While the restraint on the DBC limits the ligand exploration to poses close to the crystal bound state, a significant proportion of peripheral poses are visited during REST2 sampling. The inspection of trajectories revealed that these poses vary from one simulation to the other and form few specific interactions with the binding site. While they may have some biological relevance, we chose a strict definition for the bound state within its main free-energy well. To help us individuate adequate criteria, we define another collective variable complementary to the DBC (*c*): the sum of the distances between the N-terminal nitrogen (Phe1-N) and each of its H-bound mannoside oxygens, O-2 and O-6, respectively noted *d*_N,O2_ and *d*_N,O6_. Projecting the REST2 trajectories on the 2-D space of *δ* and *d*_N,O2_ + *d*_N,O6_ allows robust identification of an energy basin and the final criterion for the canonical bound state is expressed with the indicator function 𝟙:

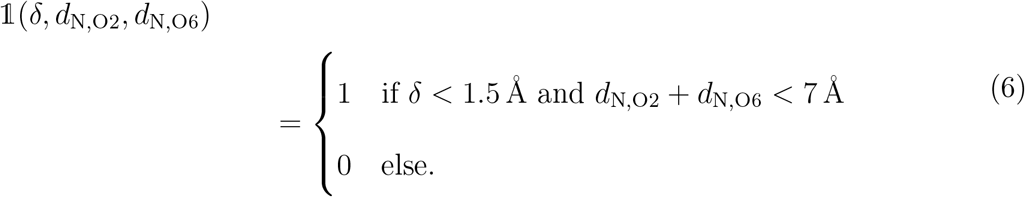

In analyzes for *bound* systems, we simply discarded all frames where the criterion is not met.

### Opening coordinate

In order to define the opening coordinate *ω*, we select a set of *n*_*ω*_ *α*-carbons belonging to the clamp loop and pocket zipper (resid. 1–16) as well as the beta-sandwich structure (resid. 17–23, 35–46, 54–58, 74–110, 126–135, 143–150), and excluding the interdomain region, beta-bulge, alpha-switch and L2/L3 flexible loops bordering the binding site. The atomic coordinate vector is linearly projected on the displacement vector between two reference structures in the *S*_*cl*_ (PDB: 4xoc) and *A*_*op*_ (PDB: 4xod) states. Let **x**_*cl*_ and **x**_*op*_ denote the 3 × *n*_*ω*_ reference coordinate vectors. Let the transformation *T*_*cl*_(**x**) denote the optimal rototranslational alignment of **x** to **x**_*cl*_. We write Δ**x**_c*l*→*op*_ = *T*_*cl*_(**x**_*op*_) *-* **x**_*cl*_ the opening displacement vector. Similarly, we can write the displacement vector with respect to **x**_*cl*_ at time *t* : Δ**x**_*cl*→*t*_ = *T*_*cl*_(**x**_*t*_) *-* **x**_*cl*_. The opening coordinate at time *t* is the dot product of Δ**x**_*cl*→ *t*_ and Δ**x**_*cl*→ *op*_ :

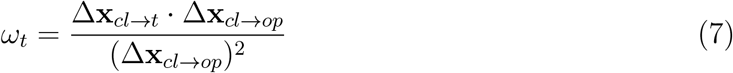

where the denominator allows to normalize between 0 (*closed*) and 1 (*open*).

### Allosteric coordinate

The allosteric coordinate *α* is built on the same principle, but include this time the *α*-carbons of the swing loop (resid. 25–34), beta bulge and alpha switch (59–71), insertion loop (111–119) and linker (152–158). The same reference structures are used to define reference coordinates **x**_*S*_ and **x**_*A*_, and similarly:

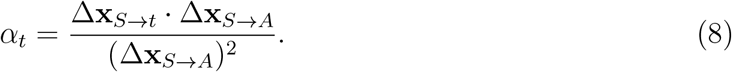

Both CVs are implemented thanks to the `PCAVARS` component of the PLUMED package. The two CVs are illustrated on Fig. 3.

**Figure 3:**
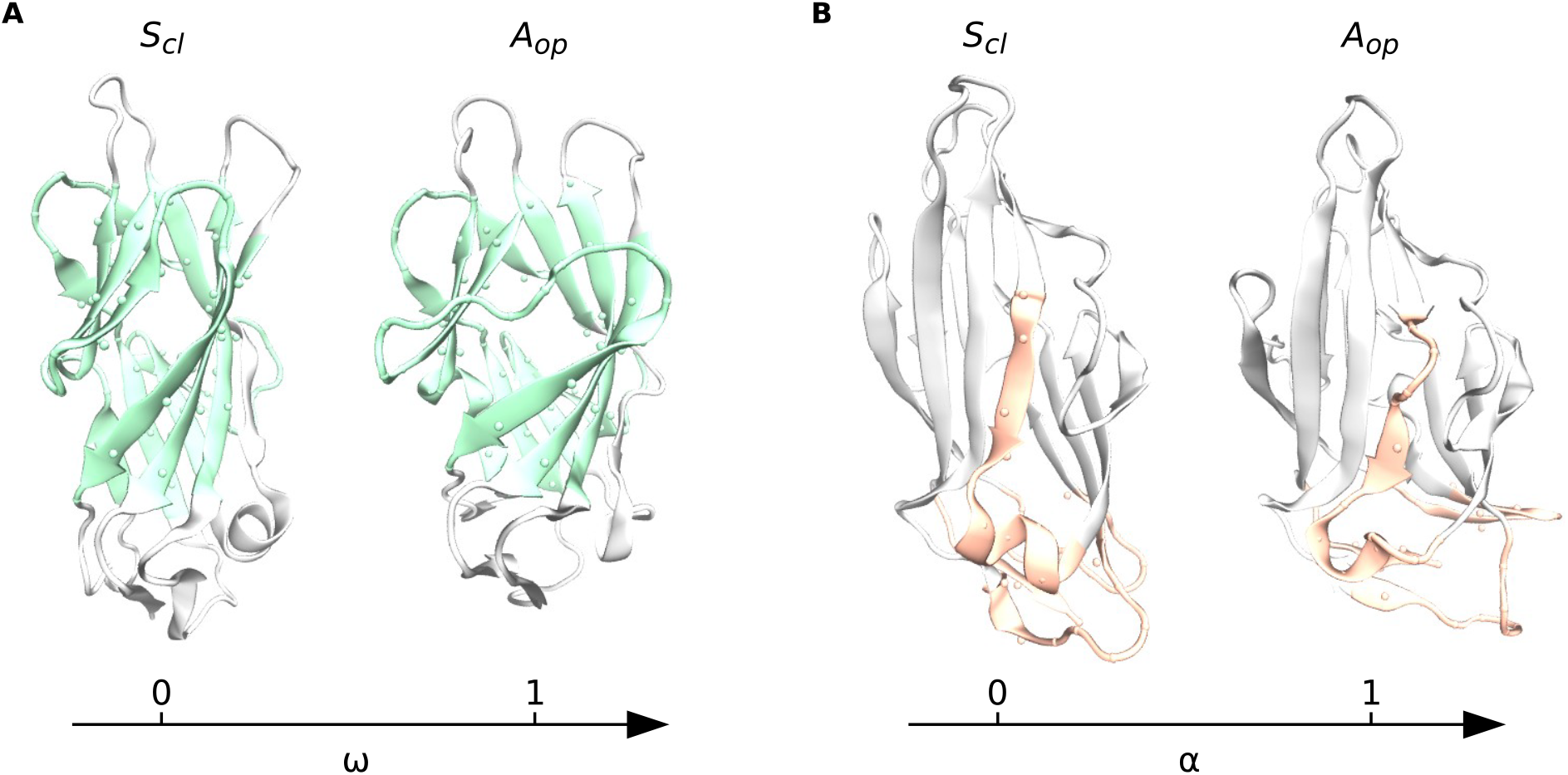
Illustration of the two collective variables used for analysis. (A) The opening coordinate *ω* is based on the beta-sandwich and clamp loop while (B) the allosteric coordinate *α* focuses on the interdomain loops and the beta-bulge/alpha-switch segment. Alpha carbons entering the CV definitions are shown as small balls and the corresponding residues are colored.

### Replica Exchange Umbrella Sampling

Computations of one-dimensional free-energy surfaces (FES) – or free-energy profiles – along the opening CV were carried out in GROMACS patched with PLUMED. We used 24 replicas with equally spaced harmonic biases (Δ *ω* = 0.05). The harmonic constant was set to *κ* = 2000 kJ mol^*-*1^ so as to ensure good overlap between adjacent windows. Windows were initialized using the REST2 data as follows: for a window *i* centered on *ω*_*i*_, we randomly select a configuration from the REST2 trajectory (restricted to the *bound* criterion, if applicable) whose opening coordinate verifies *ω*_*i*_ *-* Δ*ω < ω < ω*_*i*_ + Δ*ω*, where Δ*ω* is the window spacing. If no such configuration is found, the frame with the closest *ω* to *ω*_*i*_ is chosen instead.

Exchanges between adjacent windows were attempted every 1 ps. Because of the steepness of the underlying FES, the direct implementation of the REUS method led to regions of poor overlap between adjacent windows, as revealed by poor exchange rates. Hence, we proceeded in a two-step Adaptive Umbrella Sampling (AUS)^58^ procedure. A first 100 ns REUS allowed to compute an approximate profile *F*_0_(*ω*). The inverted potential *-F*_0_(*ω*) was then used as an external bias for a second round of 400 ns REUS, leading to a second profile *F*_1_(*ω*). The unbiased surface *F*_*ref*_ was finally recovered as:

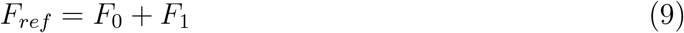

The weighted histogram analysis method (WHAM) was used to compute all free energy profiles using Grossfield‘s implementation.^59^ Statistical errors were estimated by bootstrap on 20 iterations. The quality of convergence was appreciated by comparing the profiles obtained from a two-block analysis (2 × 150 ns).

## Results and discussion

### Exploration of the conformational landscape unveils binding site plasticity

We first investigate whether the conformational landscape exploration of the lectin domain in several key states of FimH reveals the presence of protein conformations that are different from the static crystal structures. To do so, we use replica exchange with solute scaling (REST2),^55^ in which the system benefits from faster configurational exploration at higher “effective temperatures” while sampling the simulation ensemble of interest (see Methods).

We applied the REST2 sampling method to the following systems to obtain 500 ns trajectories:

1. the lectin domain in isolation, bound to the high-affinity mannose derivative *α*-D-heptylmannose (HMan), in its native *Separated* -*closed* state (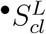, PDB: 4xoc)
2. the full FimH protein stabilized by the FimG donnor strand (DsG), bound to HMan, in the *Associated-closed* state (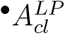, PDB: 4xoe)
3. the lectin domain in isolation as extracted from the previous structure after in-silico removal of the pilin domain (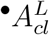, PDB: 4xoe).

Our goal here is to assess the conformational flexibility in the two identified allosteric state (*S* and *A*). We aim to investigate the binding site plasticity (in particular its propensity to open), as well as the possibility of interchange between *A* and *S* upon removal of the regulatory domain. In order to quantify the impact of the ligand on the binding site, we also simulated the three identical systems after removal of HMan. To prevent the ligand from escaping in the *bound* systems, we use a Distance-from-Bound-Configuration (DBC) collective variable, together with a flat-well restraint potential,^53^ which allows unbiased sampling of the system degrees of freedom as long as the ligand remains far away from the restraint wall (see Methods). To characterize the *open* or *closed* character of the binding site, we define an “opening” collective variable (CV) *ω*, as well as an “allosteric” or “association” variable *α* that quantifies the transition between the *S* (*α* = 0) and the *A* (*α* = 1) states (see Methods and Fig. 3).

The conformational space explored along the two CVs *ω* and *α* is shown in Fig. 4. During this exploration, all systems visit conformational states that significantly differ from the starting crystal structures. The most striking observation is perhaps that, while all simulations start in the *closed* part of the *ω*-space (*ω* ≈0), all of them spontaneously explore configurations near the *open* reference (*ω* ≈ 1). While the *A*_*op*_ state was already known for the unliganted, two-domain protein (Fig. 2), the *open* conformation for the *Separated* state *S*_*op*_ counterpart has never been observed before. However, this might not be surprising as this *open* conformation appear to be much less populated than the *closed* conformation, both in the apo and holo states.

**Figure 4:**
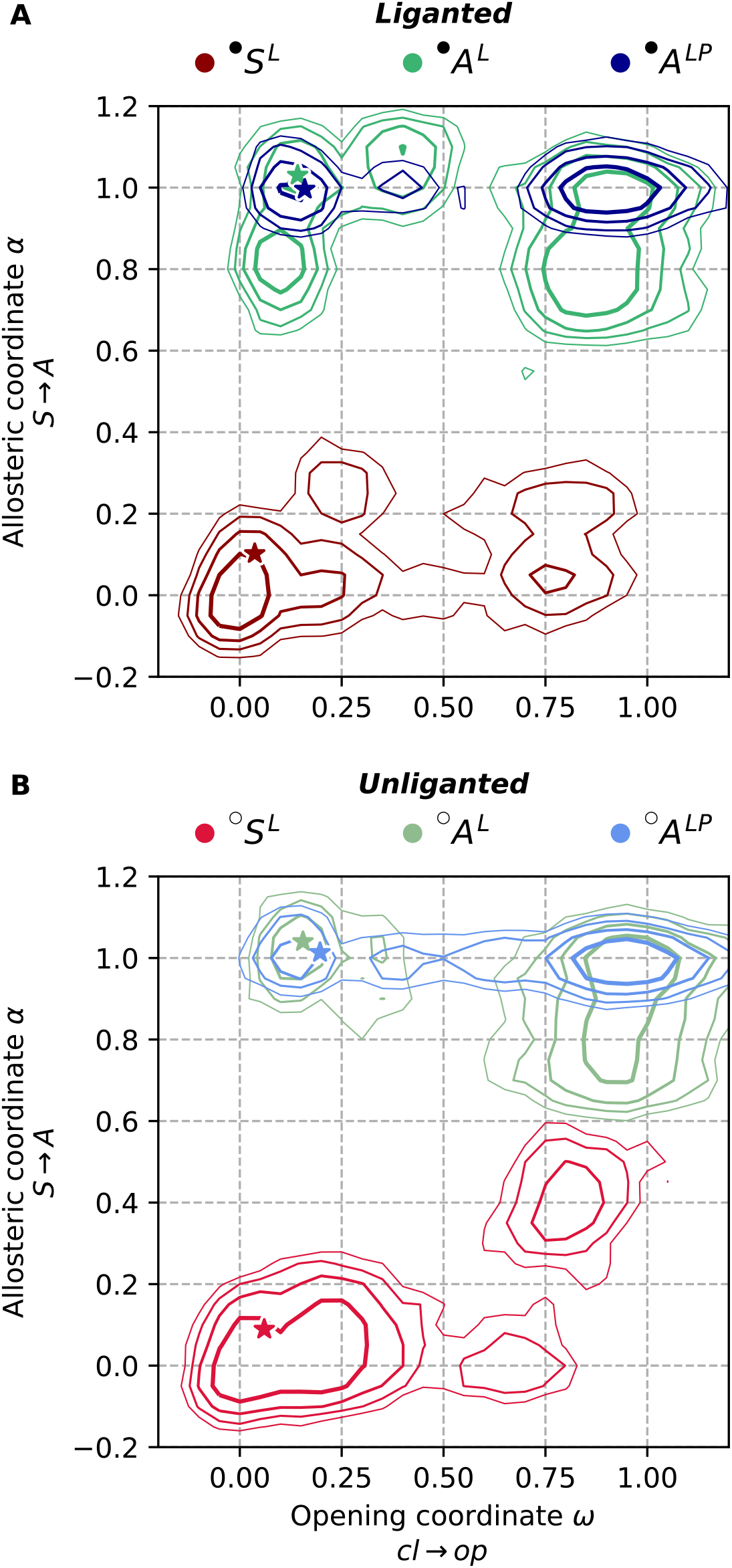
Conformational space sampled by the different REST2 simulation setups, projected on the allosteric (*S* to *A*) and opening (*cl* to *op*) collective variables, for (A) liganted and unliganted systems. Densities are estimated using a Gaussian kernel density estimate (bandwidth = 0.02) and shown as a contour plot. Stars indicate the starting configurations for each simulation.

Additionally, Fig. 4 reveals the presence of intermediate density peaks. The analysis of marginal distributions along the *ω* coordinate (see SI) allows to identify three peaks corresponding to different opening states that we will refer to as *open* (*op*), *semi-open* (*so*) and *closed* (*cl*) in the following. Snapshots of representative configurations are available in the SI.

We pause here to highlight the efficiency of the REST2 method: Indeed, preliminary microsecond MD simulations without any bias or enhanced sampling did not reveal such conformational heterogeneity. In particular, a full transition from *A*_*cl*_ to *A*_*op*_ was not observed, though partial opening to a *semi-open* state occurred (see SI). Additionally, the *S*_*cl*_ state was stable on that timescale, and no other conformations were observed. This illustrates the interest of using such unbiased enhanced-sampling techniques to provide a more comprehensive picture of the accessible conformational space, which is otherwise difficult to attain, even in relatively “long” unperturbed simulations.

However, while our approach allows to cross barriers that are on the order of 10– 20 kJ mol^*-*1^ (as we will properly estimate in the following section) on a sub-microsecond timescale, larger barriers remain inaccessible. In particular, we did not observe any transition from the *A* to the *S* configuration after artificial removal of the pilin domain (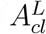 starting state), as we initially hoped. Some degree of flexibility along the *α* coordinate is observed for the lectin-only systems, though, as shown by density peaks centered on intermediate values. Visual inspection of the trajectories indicates that these are due to substantial flexibility of the interdomain region comprising the insertion loop, swing loop and linker. Nevertheless, the beta-bulge and alpha-switch motifs show remarkable stability and their reorganization probably constitutes the main kinetic obstacle for the *A* ↔ *S* transition. Less surprisingly, the full-protein *A*^*LP*^ systems display very little flexibility along the allosteric coordinate *α* because stable interactions with the pilin domain lock the lectin-domain interdomain region in its *Associated* geometry.

While the Metropolis exchange scheme undermines the system’s kinetics, REST2 sampling remains sensitive to the timescale separation between processes that exhibit significant differences in free-energy barriers. We can thus conclude that the binding-site opening dynamics are significantly faster than allosteric relaxation. This is consistent with the fact that site opening mostly involves hydrogen-bond breaking in the “zipper” region and elastic deformation of the beta-sandwich fold, while the *A* ↔ *S* transition requires a highly nonlinear reorganization of the beta-bulge/alpha-switch segment, with 180-degree flipping of several aminoacids such as Ser62 and Tyr64.^22^

Because *A*^*L*^ does not become *S*^*L*^ within the duration of REST2 simulations, we can assume that the obtained conformational ensembles are far from global equilibrium, although they might approach local equilibria within the *A* and *S* basins respectively. The robust metastability of the isolated, *A*-state lectin domain is also supported by the existence of a crystal structure of such conformation in a single-mutation variant (R60P, PDB: 5mca).^60^ However, despite the relatively long-lived nature of the *A*^*L*^ state in silico, there is little evidence that wild-type variants exhibit a significant population of *A* when the domains are separated and the linker is extended.

In this first exploratory phase, we have predicted new conformations yet unseen in experimental structures. Despite being qualitative, these findings may indicate a more nuanced picture than the two-state one suggested by previous publications. However, in addition to possible force-field biases, REST2 trajectories cannot provide a reliable estimation of the relative stabilities of the observed opening states. Indeed, the REST2 strategy is unbiased, in the sense that the system is free to explore any accessible region of the phase space. In principle, the thermodynamics of the biomolecule along any collective variable of interest could then be obtained by adequate projection of the multidimensional free energy surface (FES). However, for a large system, the system cannot sample all possible biomolecular conformations according to their Boltzmann weights, because of the high dimensionality of the phase space results in a scarcity of data in high-energy regions. Biased methods allow to focus the computational effort along a given CV of interest, perhaps at the expense of less relevant transverse degrees of freedom. We therefore now discuss the results of such targeted free-energy estimation.

### The three opening states of FimH binding site are regulated by the pilin domain

In an attempt to precise the relative abundances of various opening states of the lectin domain in both *A* and *S* allosteric states, we performed Replica-Exchange Umbrella Sampling (REUS) simulations. REUS is a variant of the well-known Umbrella Sampling technique for estimating the FES along a chosen CV. Instead of sampling independently *N* harmonically restrained windows along the biasing coordinate, they are run in parallel and allowed to exchange configurations using a Metropolis-Hastings scheme. The method has been shown to improve convergence by enhancing relaxation of transverse degrees of freedom. ^61,62^ To initiate the procedure, we benefit from the prior exploration of the REST2 simulations by randomly choosing configurations for each window with an appropriate *ω* value, and we then proceed in a two-step, adaptive fashion, as described in the Methods section.

The results are shown on Fig. 5. In agreement with crystallographic evidence, the liganted system in its *Separated* state (•*S*^*L*^) exhibits a global minimum in the region corresponding to its *closed* form. Similarly, unliganted °*A*^*L*^ and °*A*^*LP*^ systems present global minima in their *open* conformation.

**Figure 5:**
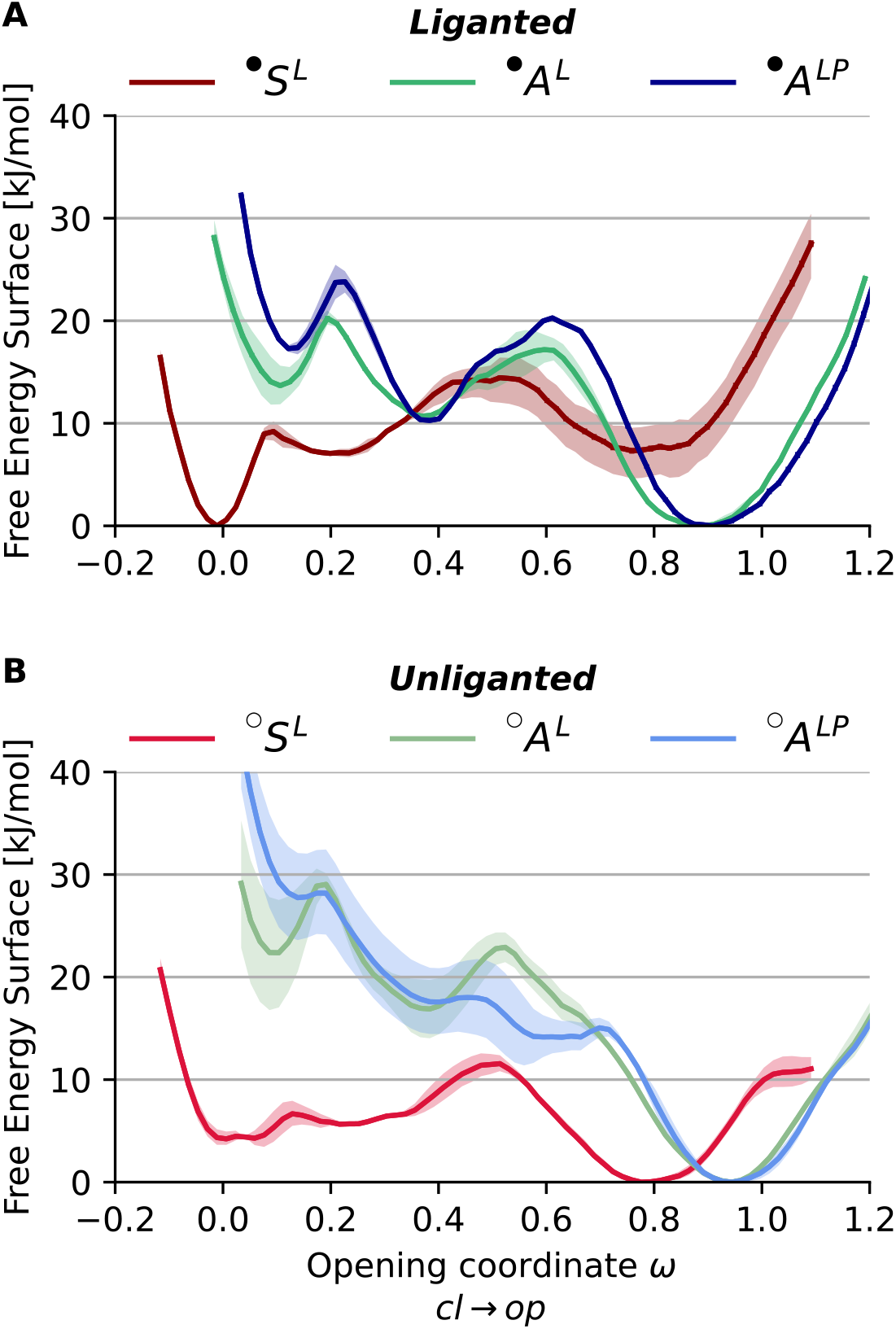
Free energy profiles along the opening coordinate (*ω*) for (A) liganted and (B) unliganted systems. Error bars indicate statistical uncertainty estimated by bootstrap (hardly visible, of the order of the plot linewidth). The shaded areas illustrate the free energy profile difference between the first and the last half of the trajectory.

In contrast, the other systems show free-energy minima that differ from their respective native crystal structures:

1. °*S*^*L*^ has a predicted minimum in its *op* state, a yet unknown conformation in crystallographic data
2. •*A*^*L*^ and •*A*^*LP*^ show a dominant *op* state, while they are *cl* in co-crystals with HMan.

The corresponding populations can be estimated by integrating the FESs between the local maxima (*ω* = 0.1; 0.5 for *S*, and *ω* = 0.2; 0.6 for *A*), as summarized in Table 1. These results suggest some conformational plasticity, at equilibrium, for both allosteric states. In particular, we predict that •*S*^*L*^ significantly populates (≈25 %) *open* and *semi-open* states in addition to the *closed* one. Even more surprisingly, the unliganted couterpart °*S*^*L*^ exhibits the reverse trend, with predominant *open* geometries (≈79 %). A similar population shift is observed for *A*^*L*^ and *A*^*LP*^ systems for which *cl* states are increased by one order of magnitude upon ligand binding. These observations provide some support to an induced-fit mechanism, where complexation prompts a conformational change in the binding site that better accommodates the ligand’s geometry. However, they invite to nuance the binary picture established from the apo and holo crystals since the *closed* binding site adjusted to the ligand is not predicted to predominate in •*A*^*LP*^.

**Table 1:** Populations of opening states, computed by Boltzmann integration of the FES (Fig. 5) for each state, as delimited by local maxima (*ω* = 0.1; 0.5 for *S*, and *ω* = 0.2; 0.6 for *A*).

| State | $P(cl)$ [%] | $P(so)$ [%] | $P(op)$ [%] |
| --- | --- | --- | --- |
| $\bullet S^L$ | 75.5 | 12.0 | 12.5 |
| $\bullet A^L$ | 0.2 | 1.1 | 98.7 |
| $\bullet A^{LP}$ | $5 \times 10^{-2}$ | 1.1 | 98.9 |
| ${}^\circ S^L$ | 11.5 | 9.6 | 78.9 |
| ${}^\circ A^L$ | $7 \times 10^{-3}$ | 0.1 | 99.9 |
| ${}^\circ A^{LP}$ | $8 \times 10^{-4}$ | 0.5 | 99.5 |

The fact that the •*A*^*LP*^ allosteric state is more stable with an *open* clamp loop configuration may offer an explanation to the vastly different unbinding kinetics measured in experiments. The lectin-only construct was shown to exhibit a *k*_off_ up to five orders of magnitude slower than the FimH·DsG one, which has been widely interpreted through the lens of unique underlying conformers, with the intriguing result that the 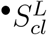 and 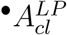 states have drastically different unbinding kinetics despite near-identical binding-site geometries in the X-ray structures. This paradox might be resolved if the main state is not the *closed* one as suggested by crystal structures, or if partial or total opening of the binding site prior to unbinding is the most probable route. It has also been suggested that despite their structural similarity, the two states exhibit different dynamics in the clamp loop region, •*A*_*cl*_ being more flexible, which appeals to the notion of dynamic allostery.^23^ These two perspectives are similar, though our interpretation better fits conventional allostery, where the allosteric transition affects the respective populations of multiple conformational states rather than the internal dynamics of a single state.

We finally note that the *closed* state is slightly more stabilized relatively to the *open* one upon removal of the pilin domain (contrast *A*^*L*^ and *A*^*LP*^ in Fig. 5). This might be related to an enhanced flexibility in the interdomain region when the pilin domain is not present (as can be seen by the wider exploration along *α* for *L* systems, Fig. 4), and suggests a weak coupling between the binding site and the interdomain loops *before* any reorganization of the beta-bulge/alpha-switch segment. One might intuitively grasp this effect since the pilin domain interacts with the swing loop, which is closely connected to the clamp loop via an intermediate beta strand (see Fig. 2): Hence, removal of the pilin domain might release some steric tension in the interdomain region that is transmitted to the binding site through the protein structure.

### External force modulates binding site opening

The current consensus on the FimH force-activated allosteric mechanism is that binding affinity is modulated by the separation of the pilin and lectin domains, that triggers a transition from the *A* to the *S* state. However, as we have just seen, each of these states exhibit significant conformational fluctuations and we cannot neglect the effect of mechanical strain on the relative stabilities of these conformations. Modeling the precise effect of a hydrodynamic flow would require to deploy a specific simulation machinery,^63^ which goes beyond the scope of the current work and which remains very challenging in an all-atom setup.^36^ Still, the effect of the traction exerted by a flowing fluid onto the bacterium and then transmitted via the pilus rod to the FimH-glycan complex can be reasonably modeled as a constant force acting between the base of the lectin domain and the ligand. This situation also echoes the setup of force-spectroscopy experiments.

We analyze the effect of a tensile force on the free energy profile along the opening coordinate of two bound systems: •*S*^*L*^ and •*A*^*LP*^. A linear bias is applied to the distance *z* between the terminal carbon of the heptyl tail of HMan (C-13) and the C-terminal carbon of the protein (Thr158 for the *L* system, Lys14 of DsG for the *LP* system). We choose *F* = 30 kJ mol^*-*1^ nm^*-*1^ ≈ 50 pN, an intermediate value where the catch-bond properties start to become apparent in vivo.^26^ We re-use the first round of our adaptive umbrella sampling approach as a starting point to a 400 ns second round in the presence of force, from which we discard the first 100 ns as equilibration.

The resulting profiles under force (Fig. 6) display a significant shift with respect to the unperturbed ones, with a force-induced stabilization of the *closed* state with respect to the *open* one. For the *S*^*L*^ system, this has for main effect to divide the population of *open* states by 3 (Table 2). For the •*A*^*LP*^ system, it increases the population of *closed* state by one order of magnitude.

**Table 2:** Population shift of opening states under force, computed by Boltzmann integration of the FES (Fig. 6) for each state, as delimited by local maxima (*ω* = 0.1; 0.5 for *S*, and *ω* = 0.2; 0.6 for *A*).

| State | $F$ [pN] | $P(cl)$ [%] | $P(so)$ [%] | $P(op)$ [%] |
| --- | --- | --- | --- | --- |
| $\bullet S^L$ | 0 | 75.5 | 12.0 | 12.5 |
| $\bullet S^L$ | 50 | 82.3 | 13.6 | 4.1 |
| $\bullet A^{LP}$ | 0 | $4.68 \times 10^{-4}$ | 1.06 | 98.9 |
| $\bullet A^{LP}$ | 50 | $5.01 \times 10^{-3}$ | 2.15 | 97.4 |

**Figure 6:**
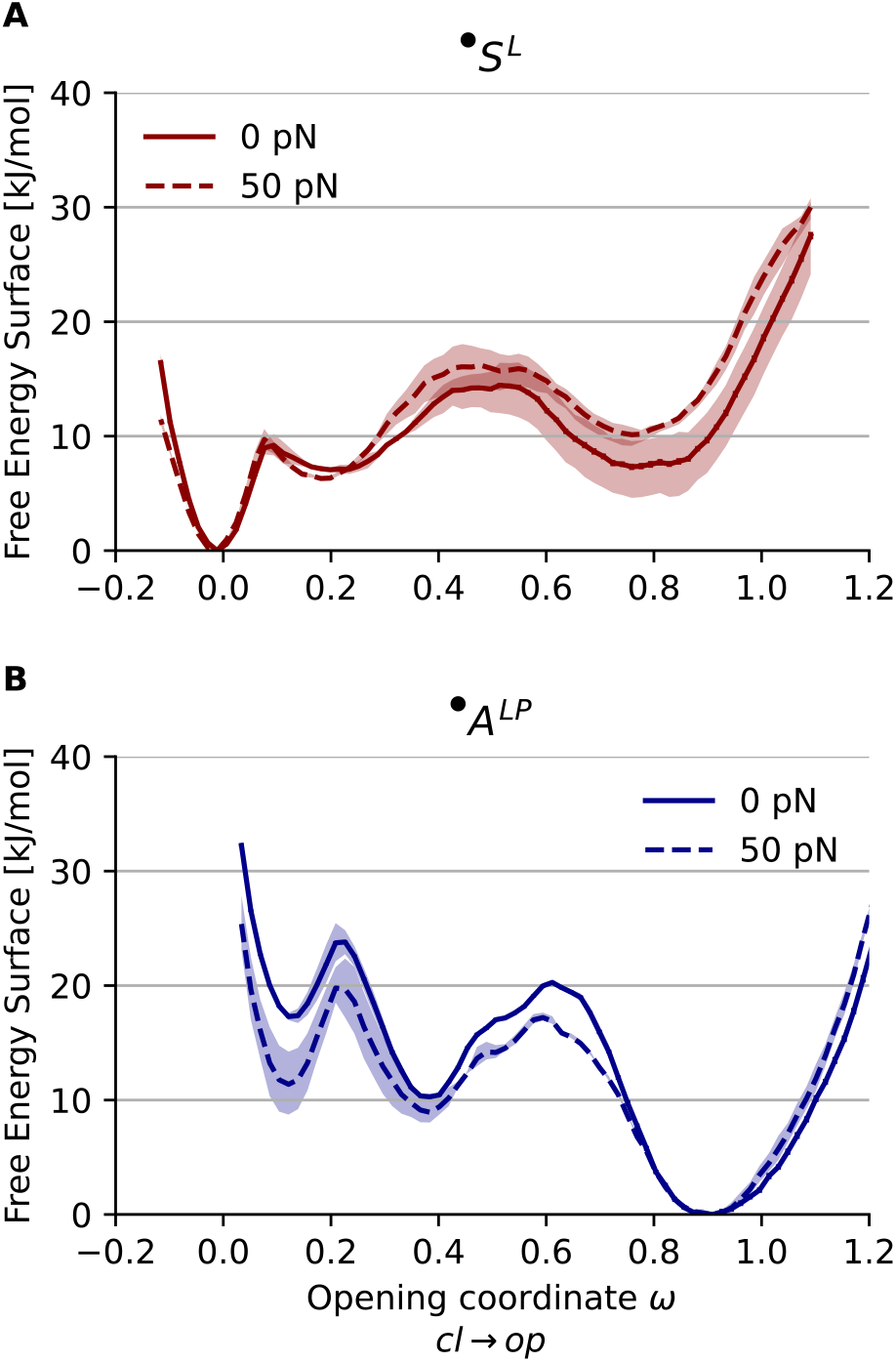
Free energy profiles along the opening coordinate (*ω*) for the liganted (A) •*S*^*L*^ and (B) •*A*^*LP*^ systems under force. Error bars indicate statistical uncertainty estimated by bootstrap (hardly visible, of the order of the plot linewidth). The shaded areas illustrate the free energy profile difference between the first and the last half of the trajectory.

While such population shifts might seem modest they could nevertheless have significant effect on the unbinding kinetics. The stabilization of the *closed* binding site conformation, which allows to retain interactions with the ligand, may thus act as a “local” catch bond effect that increases bond duration prior to domain separation and allosteric transition. Since the binding site conformational dynamics are arguably much faster than the allosteric *A* to *S* transition, as our simulations suggest and as expected from the required structural reorganizations for both processes, such a mechanism could reinforce the overall catch-bond mechanism by allowing a quick response to sudden change in mechanical strain. By reducing the relative stability of the *open* state, tensile force may thus delay unbinding and give time to the slower degrees of freedom (domain separation and *A* ↔ *S* pathway) to relax towards their most favored conformations under these conditions.

### Binding site opening influences binding affinity

We finally show that the free-energy profiles along the opening coordinate enable to obtain meaningful estimates of relative changes in the binding free-energies. Ideally, direct measurements of the binding free energies could be obtained by methods such as alchemical free energy perturbation techniques, whereby the ligand is transformed or removed from the binding site using unphysical, “alchemical” paths. Despite many attempts to estimate binding free-energies of HMan to the lectin domain in several of its relevant conformations, such calculations appeared very delicate to converge and error bars typically exceeded the relative differences between the experimental binding affinities.

However, the comparison between liganted and unliganted free-energy profiles allows to extract relative binding free-energy differences between the three opening states for a given protein system. Indeed, the relative binding free-energy change between two opening states *X* and *Y* can be expressed as

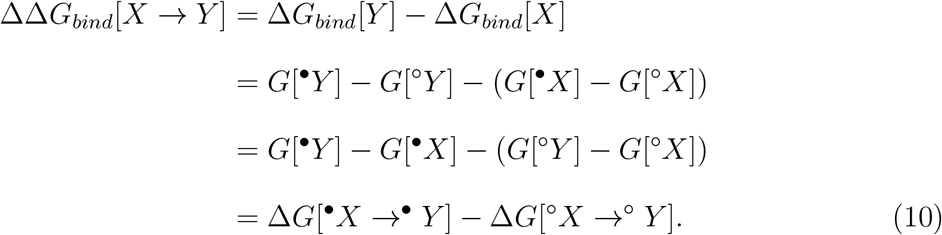

The free-energy differences in the bound (Δ*G*[•*X* → • *Y*]) and unbound (Δ*G*[•*X* → ° *Y*]) states are directly accessible from the populations computed in Table 1, using Δ*G*[*X* → *Y*] = *-k*_*B*_*T* log[*P* (*Y*)*/P* (*X*)]. Note however that we cannot compare between the *S*^*L*^, *A*^*L*^ and *A*^*LP*^ states, because free-energy profiles are defined up to an arbitrary constant. The results are expressed in Table 3, taking for each system the *cl* state as the reference.

**Table 3:** Relative binding free energies between opening states, computed for each system using Eq. 10, taking the closed state as reference.

| State | $\Delta\Delta G_{bind}[cl \rightarrow X]$ [kJ mol<sup>-1</sup>] | | |
| --- | --- | --- | --- |
|  | *cl* | *so* | *op* |
| $S^L$ | 0 | 4.2 | 9.3 |
| $A^L$ | 0 | 2.7 | 8.6 |
| $A^{LP}$ | 0 | 8.3 | 10.3 |

As anticipated, the *open* states show a lower binding affinity for all systems, with a binding free energy about 10 kJ mol^*-*1^ higher than the *closed* ones. Intermediate *semi-open* states correspond to intermediate affinities. This is overall consistent with the expectation that the *closed* binding site exhibits stronger affinity for the ligand, in part because of favorable hydrophobic interactions between the pyranose cycle and the clamp loop’s upper ridge (Ile13, Gly14) and reduced exposure to the solvent.^64^

We stress again that we cannot directly compare the *S* and *A* states. However, given the very similar binding sites of the corresponding X-ray structures, it would be tempting to attribute, at least partially, the experimentally-observed 20 kJ mol^*-*1^ difference in binding free energy^23,32^ to the difference in binding site opening, that we estimate here to contribute up to a 10 kJ mol^*-*1^ stabilization of the *bound* conformation as compared to the *unbound* one. The remaining stabilization of the bound-state in the *S* form could be due to different local fluctuations and entropic effects that may significantly influence binding.

## Conclusion

The current consensus on the FimH catch-bond mechanism is that it possesses two main allosteric states *A* (low affinity) and *S* (high affinity), with a *A* to *S* transition triggered by the pilin domain separation from the lectin (binding) domain upon application of a mechanical force. In this article, we revisit this picture by focusing on the conformational plasticity of the FimH lectin domain, using enhanced-sampling methods that enable extensive exploration of the protein free-energy landscape. We show that such considerations are essential to understand the differences in binding affinity between the *A* and *S* states, as well as previously unappreciated *intrinsic* effects of force on the binding site.

So far, most computational approaches have employed traditional MD simulations with trajectories rarely exceeding a few hundred nanoseconds. ^23,27,30,40–44^ While they provided in-sighful molecular support to experimental measurements, some important questions remain unanswered. In particular, the crystal structures of the two ligand-bound allosteric states display superimposable binding-site geometries. Previous computational work identified different fluctuation amplitudes in this region for the *S* and *A* states,^23,27,43^ and the increased flexibility in the *A* state was proposed as an explanation for the binding free-energy difference (which is about 20 kJ mol^*-*1^).^23^ In another work, Interlandi and Thomas ^64^ noticed that the loop flexibility seen in the *A*-state simulations was much higher than expected from crystallographic B-factors, suggesting possible artifacts due to crystal packing or refinement procedures.

By deploying here a combination of enhanced-sampling strategies on all-atom MD simulations with state-of-the-art protein force fields, we can provide a richer picture of the lectin domain’s conformational landscape than that offered by crystal structures alone. For each allosteric state, both in the liganted and unliganted cases, we identify three binding-site opening substates and we quantify the free-energy differences and barriers between them. This approach leads to two important conclusions. First, the comparison between liganted and unliganted systems suggests that, upon ligand binding, *closed* site conformations are always stabilized by about 10 kJ mol^*-*1^ with respect to *open* ones, supporting the idea of an induced-fit mechanism. Second, the most stable conformation for the liganted *A* state corresponds to an *open* site, in contrast to the native cocrystal structure, which is *closed*.

As we show, the experimentally-observed difference in binding affinity for the *LP* and *L* systems is actually very much compatible with the most stable site conformation being respectively *open* and *closed*. This both provides indirect support for the (a priori) surprising disagreement between the known crystal structure of •*A*^*LP*^ and our predictions, and offers a more quantitative interpretation of binding affinities than arguments based on distinct dynamics of similar site structures. We cannot completely rule out that force-field or methodological biases have over-favored *open* states in silico. On the other hand, X-ray crystallography is also vulnerable to the influence of crystallization parameters (p*H*, temperature) and crystal packing artifacts. Advanced NMR experiments could help scrutinize the opening dynamics in vitro and complement simulation data through integrative modeling.

Our approach also allows to quantify the effect of mechanical force on this conformational equilibrium. Typically, force is known to trigger the pilin-lectin domain separation, which prompts *A* to *S* transition. Our results show that after the allosteric shift, the initial 15 kJ mol^*-*1^ destabilization of the *closed* state with respect to the *open* one becomes a stabilization of about *-*10 kJ mol^*-*1^, resulting in an overall ≈*-* 25 kJ mol^*-*1^ relative stabilization. However, we also demonstrate that force has a small but previously unappreciated effect on the conformational equilibrium of the binding site in both the *A* and *S* states themselves. A pulling force in the 50 pN range (at which the catch-bond properties start to become apparent in vivo^26^) result in a 4–7 kJ mol^*-*1^ additional relative stabilization of the *closed* conformation. This phenomenon may thus act as a “local” catch-bond effect that could help the system adapt to rapid changes in tensile force and provide a temporary increase in binding strength in the *A* state before the slower allosteric transition happens along with reinforcing the bond once the transition toward the *S* state is completed.

We argue that such a two-scale catch-bond mechanism is actually relevant from an evolutionary perspective. Indeed, one may wonder how the intricate allosteric transition suggested by *A* and *S* states evolved, since natural selection can only select for an already-existing feature, and such a complex reorganization seems improbable to appear accidentally. However, two-state allosteric models are by no means necessary to produce catch-bond behaviors, and “simpler” structural models have been proposed, including deformation models, where elastic deformation of one or both partners increases favorable binding interactions. Dansuk and Keten proposed an example of such idealized, tweezer-like model, that could represent FimH’s binding site in its various opening states.^65^ This kind of very simple model with few degrees of freedom show that protein do not need to evolve complex allosteric transition pathways (such as the *A* ↔ *S* one) to acquire catch-bond properties. Hence, we may hypothesize that the local, tweezer-like catch-bond mechanism evolved first from a non-catch-bond (or *slip-bond*) receptor, and served as a basis to the more sophisticated interdomain regulation. Overall, our work illustrates the power of enhanced-sampling strategies in all-atom molecular dynamics simulation of biomolecular systems exhibiting complex interactions, providing new perspectives that cannot be easily addressed in experimental studies of the FimH catch-bond system. This kind of approaches could be beneficial to the investigation of other catch-bond systems notably involved in mechanosensing and mechanotransduction processes.^39^

## Author Contributions

O.L.-C., F.S. and G.S. conceived research. O.L.-C. performed and analyzed simulations; all authors wrote the paper and contributed in revising and editing the manuscript.

## Acknowlegment

This work was supported by the “Initiative d’Excellence” program from the French State (Grant “DYNAMO”, ANR-11-LABX-0011-01 to G.S. and F.S.), and by a PhD Fellowship “CDSN” awarded by the Ministry of National Education to O.L.-C. The simulations presented here benefited from a local computing platform administered by G. Letessier, and benefited from the HPC resources of TGCC under the allocations A0070811005 and A0090811005 made by GENCI (Grand Equipement National de Calcul Intensif).

## References

(1) Dembo, M.; Torney, D. C.; Saxman, K.; Hammer, D. The reaction-limited kinetics of membrane-to-surface adhesion and detachment. Proc. R. Soc. Lond. B. 1988, 234, 55–83.

(2) Thomas, W. E.; Vogel, V.; Sokurenko, E. Biophysics of Catch Bonds. Annu. Rev. Biophys. 2008, 37, 399–416.

(3) Sokurenko, E. V.; Vogel, V.; Thomas, W. E. Catch-Bond Mechanism of Force-Enhanced Adhesion: Counterintuitive, Elusive, but … Widespread? Cell Host Microbe 2008, 4, 314–323.

(4) Marshall, B. T.; Long, M.; Piper, J. W.; Yago, T.; McEver, R. P.; Zhu, C. Direct observation of catch bonds involving cell-adhesion molecules. Nature 2003, 423, 190– 193.

(5) Yago, T.; Lou, J.; Wu, T.; Yang, J.; Miner, J. J.; Coburn, L.; López, J. A.; Cruz, M. A.; Dong, J.-F.; McIntire, L. V. et al. Platelet glycoprotein Iba forms catch bonds with human WT vWF but not with type 2B von Willebrand disease vWF. J. Clin. Invest. 2008, 118, 3195–3207.

(6) Huang, D. L.; Bax, N. A.; Buckley, C. D.; Weis, W. I.; Dunn, A. R. Vinculin forms a directionally asymmetric catch bond with F-actin. Science 2017, 357, 703–706.

(7) Vitry, P.; Valotteau, C.; Feuillie, C.; Bernard, S.; Alsteens, D.; Geoghegan, J. A.; Dufrêne, Y. F. Force-Induced Strengthening of the Interaction between Staphylococcus aureus Clumping Factor B and Loricrin. mBio 2017, 8, e01748–17.

(8) Herman-Bausier, P.; Labate, C.; Towell, A. M.; Derclaye, S.; Geoghegan, J. A.; Dufrêne, Y. F. Staphylococcus aureus clumping factor A is a force-sensitive molecular switch that activates bacterial adhesion. Proc. Natl. Acad. Sci. U.S.A. 2018, 115, 5564–5569.

(9) Xu, X.-P.; Pokutta, S.; Torres, M.; Swift, M. F.; Hanein, D.; Volkmann, N.; Weis, W. I. Structural basis of aE-catenin–F-actin catch bond behavior. eLife 2020, 9, e60878.

(10) Lim, Y. B.; Thingna, J.; Kong, F.; Dao, M.; Cao, J.; Lim, C. T. Temperature-Induced Catch-Slip to Slip Bond Transit in Plasmodium falciparum-Infected Erythrocytes. Biophys. J. 2020, 118, 105–116.

(11) Liu, B.; Kolawole, E. M.; Evavold, B. D. Mechanobiology of T Cell Activation: To Catch a Bond. Annu. Rev. Cell Dev. Biol. 2021, 37, 65–87.

(12) Iskratsch, T.; Wolfenson, H.; Sheetz, M. P. Appreciating force and shape — the rise of mechanotransduction in cell biology. Nat. Rev. Mol. Cell Biol. 2014, 15, 825–833.

(13) Polacheck, W. J.; Chen, C. S. Measuring cell-generated forces: a guide to the available tools. Nat. Methods 2016, 13, 415–423.

(14) Neuman, K. C.; Nagy, A. Single-molecule force spectroscopy: optical tweezers, magnetic tweezers and atomic force microscopy. Nat. Methods 2008, 5, 491–505.

(15) Knight, S. D.; Bouckaert, J. In Glycoscience and Microbial Adhesion; Lindhorst, T. K., Oscarson, S., Eds.; Springer Berlin Heidelberg: Berlin, Heidelberg, 2009; Vol. 288; pp 67–107, Series Title: Topics in Current Chemistry.

(16) Foroogh, N.; Rezvan, M.; Ahmad, K.; Mahmood, S. Structural and functional characterization of the FimH adhesin of uropathogenic Escherichia coli and its novel applications. Microbial Pathogenesis 2021, 161, 105288.

(17) Foxman, B. Epidemiology of urinary tract infections: Incidence, morbidity, and economic costs. Dis. Mon. 2003, 49, 53–70.

(18) Terlizzi, M. E.; Gribaudo, G.; Maffei, M. E. UroPathogenic Escherichia coli (UPEC) Infections: Virulence Factors, Bladder Responses, Antibiotic, and Non-antibiotic Antimicrobial Strategies. Front. Microbiol. 2017, 8, 1566.

(19) Bouckaert, J.; Berglund, J.; Schembri, M.; De Genst, E.; Cools, L.; Wuhrer, M.; Hung, C.-S.; Pinkner, J.; Slättegård, R.; Zavialov, A. et al. Receptor binding studies disclose a novel class of high-affinity inhibitors of the Escherichia coli FimH adhesin: A novel class of FimH high-affinity ligands. Mol. Microbiol. 2004, 55, 441–455.

(20) Hartmann, M.; Lindhorst, T. K. The Bacterial Lectin FimH, a Target for Drug Discovery – Carbohydrate Inhibitors of Type 1 Fimbriae-Mediated Bacterial Adhesion. Eur. J. Org. Chem. 2011, 2011, 3583–3609.

(21) Asadi, A.; Razavi, S.; Talebi, M.; Gholami, M. A review on anti-adhesion therapies of bacterial diseases. Infection 2019, 47, 13–23.

(22) Le Trong, I.; Aprikian, P.; Kidd, B. A.; Forero-Shelton, M.; Tchesnokova, V.; Rajagopal, P.; Rodriguez, V.; Interlandi, G.; Klevit, R.; Vogel, V. et al. Structural Basis for Mechanical Force Regulation of the Adhesin FimH via Finger Trap-like b Sheet Twisting. Cell 2010, 141, 645–655.

(23) Sauer, M. M.; Jakob, R. P.; Eras, J.; Baday, S.; Eriş, D.; Navarra, G.; Bernèche, S.; Ernst, B.; Maier, T.; Glockshuber, R. Catch-bond mechanism of the bacterial adhesin FimH. Nat. Commun. 2016, 7, 10738.

(24) Thomas, W.; Forero, M.; Yakovenko, O.; Nilsson, L.; Vicini, P.; Sokurenko, E.; Vogel, V. Catch-Bond Model Derived from Allostery Explains Force-Activated Bacterial Adhesion. Biophys. J. 2006, 90, 753–764.

(25) Aprikian, P.; Tchesnokova, V.; Kidd, B.; Yakovenko, O.; Yarov-Yarovoy, V.; Trinchina, E.; Vogel, V.; Thomas, W.; Sokurenko, E. Interdomain Interaction in the FimH Adhesin of Escherichia coli Regulates the Affinity to Mannose. J. Biol. Chem. 2007, 282, 23437–23446.

(26) Yakovenko, O.; Sharma, S.; Forero, M.; Tchesnokova, V.; Aprikian, P.; Kidd, B.; Mach, A.; Vogel, V.; Sokurenko, E.; Thomas, W. E. FimH Forms Catch Bonds That Are Enhanced by Mechanical Force Due to Allosteric Regulation. J. Biol. Chem. 2008, 283, 11596–11605.

(27) Rodriguez, V. B.; Kidd, B. A.; Interlandi, G.; Tchesnokova, V.; Sokurenko, E. V.; Thomas, W. E. Allosteric Coupling in the Bacterial Adhesive Protein FimH. J. Biol. Chem. 2013, 288, 24128–24139.

(28) Kalas, V.; Pinkner, J. S.; Hannan, T. J.; Hibbing, M. E.; Dodson, K. W.; Holehouse, A. S.; Zhang, H.; Tolia, N. H.; Gross, M. L.; Pappu, R. V. et al. Evolutionary fine-tuning of conformational ensembles in FimH during host-pathogen interactions. Sci. Adv. 2017, 3, e1601944.

(29) Rabbani, S.; Fiege, B.; Eris, D.; Silbermann, M.; Jakob, R. P.; Navarra, G.; Maier, T.; Ernst, B. Conformational switch of the bacterial adhesin FimH in the absence of the regulatory domain: Engineering a minimalistic allosteric system. J. Biol. Chem. 2018, 293, 1835–1849.

(30) Alonso-Caballero, A.; Schönfelder, J.; Poly, S.; Corsetti, F.; De Sancho, D.; Artacho, E.; Perez-Jimenez, R. Mechanical architecture and folding of E. coli type 1 pilus domains. Nat. Commun. 2018, 9, 2758.

(31) Puorger, C.; Eidam, O.; Capitani, G.; Erilov, D.; Grütter, M. G.; Glockshuber, R. Infinite Kinetic Stability against Dissociation of Supramolecular Protein Complexes through Donor Strand Complementation. Structure 2008, 16, 631–642.

(32) Sauer, M. M.; Jakob, R. P.; Luber, T.; Canonica, F.; Navarra, G.; Ernst, B.; Unverzagt, C.; Maier, T.; Glockshuber, R. Binding of the Bacterial Adhesin FimH to Its Natural, Multivalent High-Mannose Type Glycan Targets. J. Am. Chem. Soc. 2019, 141, 936–944.

(33) Davis, A. M.; Teague, S. J.; Kleywegt, G. J. Application and Limitations of X-ray Crystallographic Data in Structure-Based Ligand and Drug Design. Angew. Chem. Int. Ed. 2003, 42, 2718–2736.

(34) Wlodawer, A.; Minor, W.; Dauter, Z.; Jaskolski, M. Protein crystallography for non-crystallographers, or how to get the best (but not more) from published macromolecular structures: Protein crystallography for non-crystallographers. FEBS J. 2008, 275, 1– 21.

(35) Kuzmanic, A.; Pannu, N. S.; Zagrovic, B. X-ray refinement significantly underestimates the level of microscopic heterogeneity in biomolecular crystals. Nat. Commun. 2014, 5, 3220.

(36) Stirnemann, G. Recent Advances and Emerging Challenges in the Molecular Modeling of Mechanobiological Processes. J. Phys. Chem. B 2022, 126, 1365–1374.

(37) Stirnemann, G. Molecular interpretation of single-molecule force spectroscopy experiments with computational approaches. Chem. Commun. 2022, 58, 7110–7119.

(38) Franz, F.; Daday, C.; Gräter, F. Advances in molecular simulations of protein mechanical properties and function. Curr. Opin. Struct. Biol. 2020, 61, 132–138.

(39) Gomez, D.; Peña Ccoa, W. J.; Singh, Y.; Rojas, E.; Hocky, G. M. Molecular Paradigms for Biological Mechanosensing. J. Phys. Chem. B 2021, 125, 12115–12124.

(40) Thomas, W. E.; Trintchina, E.; Forero, M.; Vogel, V.; Sokurenko, E. V. Bacterial Adhesion to Target Cells Enhanced by Shear Force. Cell 2002, 109, 913–923.

(41) Aprikian, P.; Interlandi, G.; Kidd, B. A.; Le Trong, I.; Tchesnokova, V.; Yakovenko, O.; Whitfield, M. J.; Bullitt, E.; Stenkamp, R. E.; Thomas, W. E. et al. The Bacterial Fimbrial Tip Acts as a Mechanical Force Sensor. PLoS Biol. 2011, 9, e1000617.

(42) Nilsson, L. M.; Yakovenko, O.; Tchesnokova, V.; Thomas, W. E.; Schembri, M. A.; Vogel, V.; Klemm, P.; Sokurenko, E. V. The cysteine bond in the Escherichia coli FimH adhesin is critical for adhesion under flow conditions. Mol. Microbiol. 2007, 65, 1158–1169.

(43) Singaravelu, M.; Selvan, A.; Anishetty, S. Molecular dynamics simulations of lectin domain of FimH and immunoinformatics for the design of potential vaccine candidates. Comput. Biol. Chem. 2014, 52, 18–24.

(44) Liu, J.; Amaral, L. A. N.; Keten, S. Conformational stability of the bacterial adhesin, FimH, with an inactivating mutation. Proteins 2021, 89, 276–288.

(45) Magala, P.; Klevit, R. E.; Thomas, W. E.; Sokurenko, E. V.; Stenkamp, R. E. RMSD analysis of structures of the bacterial protein FimH identifies five conformations of its lectin domain. Proteins 2020, 88, 593–603.

(46) Geibel, S.; Procko, E.; Hultgren, S. J.; Baker, D.; Waksman, G. Structural and energetic basis of folded-protein transport by the FimD usher. Nature 2013, 496, 243–246.

(47) Choudhury, D.; Thompson, A.; Stojanoff, V.; Langermann, S.; Pinkner, J.; Hultgren, S. J.; Knight, S. D. X-ray Structure of the FimC-FimH Chaperone-Adhesin Complex from Uropathogenic Escherichia coli. Science 1999, 285, 1061–1066.

(48) Abraham, M. J.; Murtola, T.; Schulz, R.; Páll, S.; Smith, J. C.; Hess, B.; Lindahl, E. GROMACS: High performance molecular simulations through multi-level parallelism from laptops to supercomputers. SoftwareX 2015, 1-2, 19–25.

(49) Lindorff-Larsen, K.; Piana, S.; Palmo, K.; Maragakis, P.; Klepeis, J. L.; Dror, R. O.; Shaw, D. E. Improved side-chain torsion potentials for the Amber ff99SB protein force field: Improved Protein Side-Chain Potentials. Proteins 2010, 78, 1950–1958.

(50) Anandakrishnan, R.; Aguilar, B.; Onufriev, A. V. H++ 3.0: automating pK prediction and the preparation of biomolecular structures for atomistic molecular modeling and simulations. Nucleic Acids Res. 2012, 40, W537–W541.

(51) Wang, J.; Wolf, R. M.; Caldwell, J. W.; Kollman, P. A.; Case, D. A. Development and testing of a general amber force field. J. Comput. Chem. 2004, 25, 1157–1174.

(52) Sousa da Silva, A. W.; Vranken, W. F. ACPYPE - AnteChamber PYthon Parser interfacE. BMC Res. Notes 2012, 5, 367.

(53) Salari, R.; Joseph, T.; Lohia, R.; Hénin, J.; Brannigan, G. A Streamlined, General Approach for Computing Ligand Binding Free Energies and Its Application to GPCR-Bound Cholesterol. J. Chem. Theory Comput. 2018, 14, 6560–6573.

(54) Fiorin, G.; Klein, M. L.; Hénin, J. Using collective variables to drive molecular dynamics simulations. Mol. Phys. 2013, 111, 3345–3362.

(55) Wang, L.; Friesner, R. A.; Berne, B. J. Replica Exchange with Solute Scaling: A More Efficient Version of Replica Exchange with Solute Tempering (REST2). J. Phys. Chem. B 2011, 115, 9431–9438.

(56) The PLUMED consortium, Promoting transparency and reproducibility in enhanced molecular simulations. Nat. Methods 2019, 16, 670–673.

(57) Tribello, G. A.; Bonomi, M.; Branduardi, D.; Camilloni, C.; Bussi, G. PLUMED 2: New feathers for an old bird. Comput. Phys. Commun. 2014, 185, 604–613.

(58) Mezei, M. Adaptive umbrella sampling: Self-consistent determination of the non-Boltzmann bias. J. Comput. Phys. 1987, 68, 237–248.

(59) Grossfield, A. WHAM: the weighted histogram analysis method. http://membrane.urmc.rochester.edu/wordpress/?page_id=126.

(60) Schwartz, D. J.; Kalas, V.; Pinkner, J. S.; Chen, S. L.; Spaulding, C. N.; Dodson, K. W.; Hultgren, S. J. Positively selected FimH residues enhance virulence during urinary tract infection by altering FimH conformation. Proc. Natl. Acad. Sci. U.S.A. 2013, 110, 15530–15537.

(61) Wingbermühle, S.; Schäfer, L. V. Capturing the Flexibility of a Protein–Ligand Complex: Binding Free Energies from Different Enhanced Sampling Techniques. J. Chem. Theory Comput. 2020, 16, 4615–4630.

(62) Kästner, J. Umbrella sampling: Umbrella sampling. WIREs Comput. Mol. Sci. 2011, 1, 932–942.

(63) Sterpone, F.; Doutreligne, S.; Tran, T. T.; Melchionna, S.; Baaden, M.; Nguyen, P. H.; Derreumaux, P. Multi-scale simulations of biological systems using the OPEP coarsegrained model. Biochem. Biophys. Res. Commun. 2018, 498, 296–304.

(64) Interlandi, G.; Thomas, W. E. Mechanism of allosteric propagation across a b-sheet structure investigated by molecular dynamics simulations. Proteins 2016, 84, 990– 1008.

(65) Dansuk, K. C.; Keten, S. A Simple Mechanical Model for Synthetic Catch Bonds. Matter 2019, 1, 911–925.

