## Supplementary Material for "Binding site plasticity regulation of the FimH catch-bond mechanism"

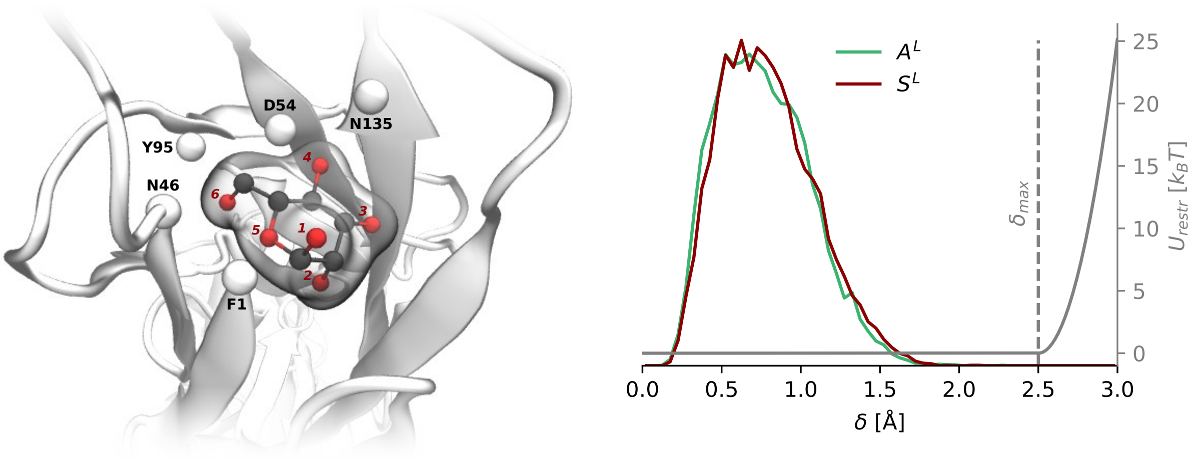


Figure S1: Choice of the DBC restraining potential width $\delta_{max}$. *(left)* Illustration of the DBC restraining potential. Alpha carbons of the binding site (white beads) are used to define the binding site frame of reference in which the mannopyranoside cycle's atoms are allowed to fluctuate within a hypervolume around their reference position. *(right)* DBC distributions in the native state are computed from preliminary vanilla MD simulations of $^{\bullet}A^L$ and $^{\bullet}S^L$ systems, and the width $\delta_{max}$ of the flat-well restraining potential is set accordingly.

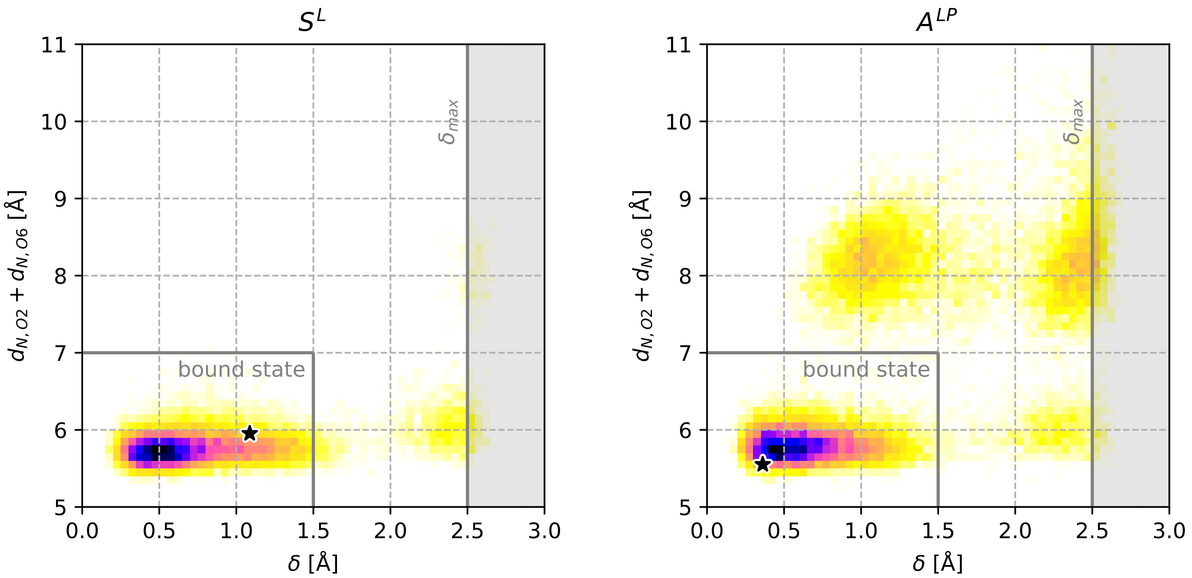


Figure S2: Definition of the bound state. While REST2 simulations explore near-bound configurations within the restraint, the joint distribution of the DBC ($\delta$) and $d_{N,O2} + d_{N,O6}$ show a well defined basin for the native *bound* state. Initial (crystal) configurations are denoted by a star ($\star$). Examples are given for $^{\bullet}S^L$ and $^{\bullet}A^{LP}$ – $^{\bullet}A^L$ show similar results.

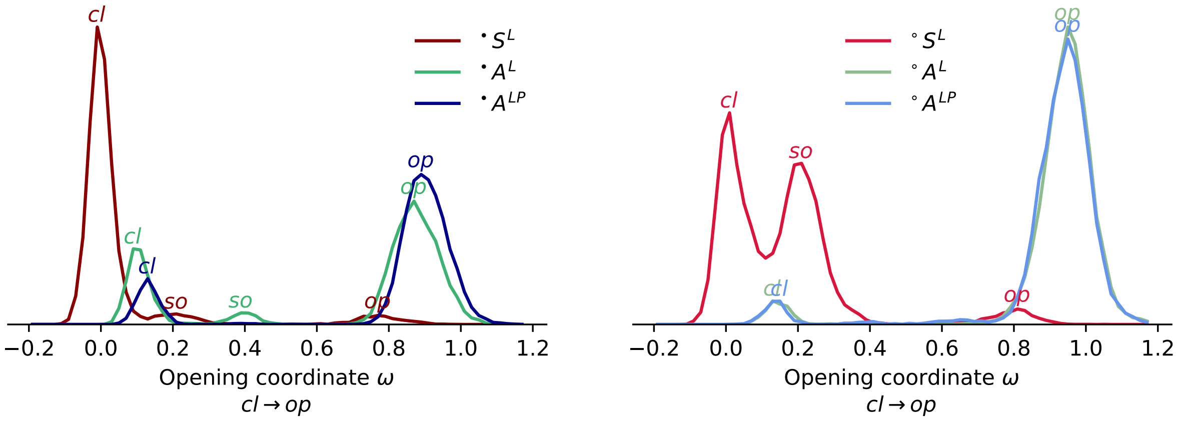


Figure S3: Marginal distributions along the opening coordinate in REST2 simulations for the liganted *(left)* and unliganted *(right)* systems. We can clearly identify peaks for the *open (op)*, *semi-open (so)* and *closed (cl)* states.

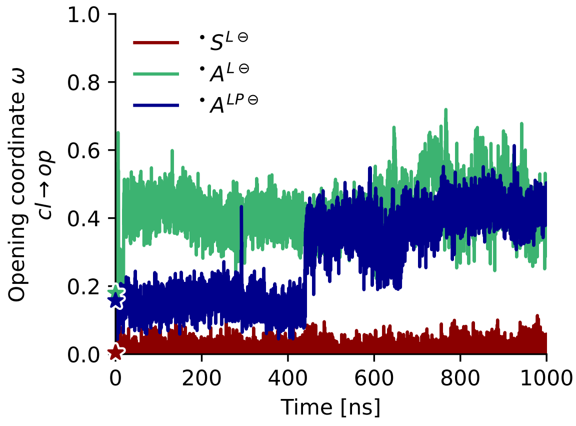


Figure S4: Evolution of the opening coordinate in microsecond-long standard MD simulations. Note that due to different system preparation, the His45 was in a monoprotonated state, as opposed to the cationic diprotonated form used in other simulations and consistent with $pK_a$ predictions. This should not have a major impact as we simply aim to illustrate here that metastability of the opening states is a limitting factor overcome by the REST2 enhanced sampling method.

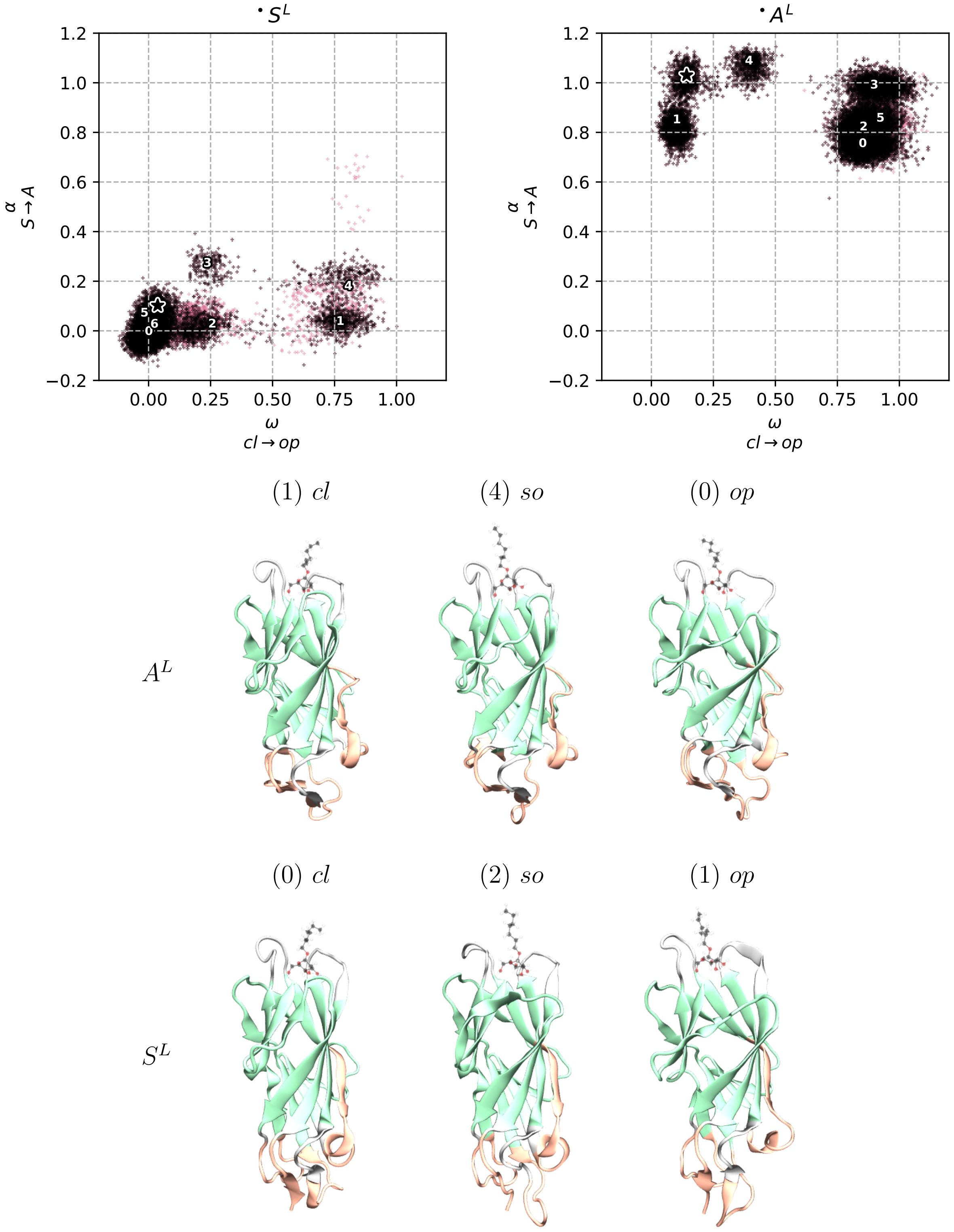


Figure S5: Representative configurations of various opening states. REST2 trajectories are projected on the ($\omega$, $\alpha$)-space, with stars denoting initial structures. Pink dots correspond to frames removed from analysis because they did not meet the bound criterion (Eq. 6). For illustrative purpose we use a clustering algorithm.[1]

### References

(1) Sittel, F.; Stock, G. Robust Density-Based Clustering To Identify Metastable Conformational States of Proteins. *J. Chem. Theory Comput.* **2016**, *12*, 2426–2435.
